# Old mice have a functionally distinct contractile reserve from obese mice

**DOI:** 10.1101/2025.05.13.653770

**Authors:** Sarah L. Sturgill, Mohammed Aidja, Benjamin Hu, Mark T. Ziolo

**Affiliations:** Dept of Physiology and Cell Biology, Davis Heart and Lung Research Institute, The Ohio State University

## Abstract

**Background:** Cardiovascular disease (CVD) is the leading cause of global mortality, with recent increases attributed to demographic shifts in age and rising rates of obesity. Diminished contractile reserve is a hallmark of a diseased heart; assessing contractile reserve is pivotal in prognosticating and monitoring CVD progression. The Frank-Starling mechanism and sympathetic stimulation are key to enhance contractile reserve but have not been explored *in vivo* in old and obese mouse models of CVD. This project aims to use speckle tracking echocardiography (STE) to characterize the function of the heart at baseline, with increased preload, and with sympathetic stimulation. We hypothesize that along with blunted systolic function, diastolic function, and contractility, old and obese mice will have a blunted contractile reserve.

**Methods:** STE was obtained for control (4- month-old), aged (24-month-old), and obese mice (high fat diet-induced). Mice received an intravenous injection of 150μL saline to increase preload to assess the Frank-Starling response, followed by injection of β1adrenergic receptor agonist dobutamine to assess sympathetic response.

**Results:** At baseline, aging and obese mice demonstrated blunted systolic, diastolic function, and contractility. Endocardial and epicardial wall displacement differed between aging and obese mice with contractile reserve, indicating that they have functionally distinct cardiac phenotypes.

**Conclusions:** This study is the first to demonstrate blunted systolic function, diastolic function, and contractility through STE in aging and obese mice. Our novel method for investigating the contractile reserve of mice demonstrated aging and obese mice have dissimilar responses when assessing contractile reserve, which could contribute to their distinct functional phenotypes.

## INTRODUCTION

Cardiovascular disease (CVD) is the leading cause of death worldwide. While great strides have been made to reduce mortality due to CVD, within the past decade, there has been an increase in mortality trends.^1^ There is a large population approaching an age (greater than 20% over the age of 65 by 2030)^2^ where CVD is the leading cause of death.^3^ Another cause of increased mortality from CVD is obesity, a comorbidity that contributes directly to cardiovascular risk factors, such as type 2 diabetes, hypertension, etc.^4^ Greater than 40% of the US population classify as obese, with an anticipated increase to 50% by 2030.^5^ Aging and obesity are risk factors for the development of heart failure with preserved ejection fraction (HFpEF)^6, 7^, with greater than ∼90% of HFpEF patients classifying as obese and/or elderly.^8, 9^

The heart’s function can be enhanced in response to increased metabolic demand from the tissues (e.g., exercise), which is defined as cardiac reserve.^10^ A major component of cardiac reserve is increased pump performance (i.e., contractile reserve), which is an increase in the contraction of the heart.^11^ A hallmark of a diseased heart is the loss of cardiac reserve caused by diminished contractile reserve, which is observed in heart failure.^12–15^ Although both aging and obesity increase the risk for HFpEF, they present as two distinct physiological phenotypes. Thus, the purpose of this study was to investigate the cardiac reserve of these two models.

Contractile reserve can be assessed *in vivo* by either increasing preload^16^ or via sympathetic stimulation.^17^ The relationship between preload (i.e., end diastolic volume) and heart performance (i.e., stroke volume) is also known as the Frank-Starling Law of the Heart. The Frank-Starling mechanism is based on the initial length of myocardial tissue and its positive relationship with force generated by contraction. An increase in preload causes the myocardial tissue to stretch, resulting in a longer initial length. This increased length causes a corresponding increase in contractile force and thus, stroke volume. Alternatively, sympathetic stimulation, another influential mechanism to access contractile reserve, occurs by activation of the β-adrenergic receptors (AR) to increase contractility. Norepinephrine, epinephrine, or pharmacological (i.e., dobutamine) activation of the receptor will induce a signaling cascade to increase heart performance via cAMP-dependent protein kinase phosphorylation of calcium-handling proteins critical for increasing contractile reserve.

Unfortunately, the Frank-Starling mechanism (the most influential mechanism to access contractile reserve) has not been investigated *in vivo* in obese or aging mouse models. Additionally, the functional effects of sympathetic stimulation *in vivo* have mostly been assessed by invasive and terminal intra left ventricular catheterization for pressure-volume (PV) measurements, such as the rate of pressure change (dP/dt) which is preload- and heart rate-dependent, or *via* MRI, which is expensive.^18, 19^ For this project, we will use speckle tracking echocardiography (STE) to measure heart function. Recent studies have highlighted the quantitative benefits of using STE to measure heart function. Studies have demonstrated that measurements of cardiac strain are less load- and heart rate-dependent than traditional catheter (i.e., dP/dt) or echo-derived measurements,^20^ which is essential for indices of contractility.^21^ STE has been demonstrated to be as determinative of heart function as highly invasive procedures like PV loops.^22^ Therefore, STE can provide alternative, non-terminal indices of systolic function (strain), diastolic function (reverse strain rate), and contractility (strain rate). We will also use traditional echocardiography to measure cardiovascular performance (defined as function of the heart and vasculature) and heart performance (defined as the overall function of the heart).^21^

Here, we also describe new techniques for modifying preload and performing sympathetic stimulation to assess contractile reserve in mice using STE. Thus, in mouse models of aging (24 months of age) and obesity (high fat diet-induced), we obtained measurements of systolic function, diastolic function, and contractility. Further, we investigated the contractile reserve (through Frank-Starling relationship and sympathetic stimulation) in these two mouse models of disease. We demonstrated that aging and obesity result in diminished systolic function, diastolic function, and contractility. Further, obesity and aging have a blunted contractile reserve.

## METHODS

All animal use protocols were approved by the Institutional Animal Care and Use Committee at The Ohio State University and meet NIH standards.

### Animal Models

We generated 4 groups of preclinical mouse models. Control mice were purchased from Charles River (male, 16-week-old C57Bl/6 wild type mice, strain 027). As a model of improved heart function, 16-week-old Charles River (male, C57Bl/6 wild type mice, strain 027) were exercised using an 8-week High Intensity Interval Training (HIIT) protocol as previously published (see Supplementary Table 7).^23, 24^ Briefly, mice underwent treadmill (Columbus Instruments, Columbus OH) HIIT for 8 weeks starting at 30 min/day and gradually increased to 80 min/day. Following a 10-minute warm up, mice were challenged at a high intensity fast pace for 4 minutes followed by 1 minute of low intensity recovery pace. This interval set was repeated until the designated time was completed and then a 4-minute cool down was provided. Experimental measurements were obtained three days after completing this 8-week HIIT program.

Two mouse models were created to recapitulate the shifting demographics of the population. Aging mice (male, C57Bl/6 WT, NIA – Charles River) were obtained at 24 months-of-age from the National Institute of Aging. For the obesity phenotype, 16-week-old male mice were put on a High Fat Diet (HFD) of 60 kcal% fat (Research Diets Inc., New Brunswick, NJ) for 16 weeks^25, 26^. HFD mice had significantly higher body weights compared to control mice (24.7 ± 0.5 g vs. 50.3 ± 1.7 g, p=<0.0001 unpaired t test).

All mouse groups were Charles River C57Bl/6 WT male background to coincide with NIA mouse background.

### Echocardiography

Mice were induced at 3-5% isoflurane with 100% O2 flow of 1.5 L/min. Anesthesia was maintained at 3% isoflurane, with adjustments made as needed to maintain a sufficiently deep plane of anesthesia. Body temperature was maintained at 37 °C and monitored by rectal thermometer. Echocardiography measurements were performed using a Vevo 3100 system (Visual Sonics) and a MX550D (40MHz) transducer. B mode cine loops were obtained from short-axis scans at the level of the papillary muscles. The probe position was maintained for all measurements (baseline, preload, and dobutamine echocardiograms).

### Speckle tracking analysis

All echocardiograms were analyzed using VevoLab software. Short axis B mode cine loops were first converted into Anatomical M-mode images with the line drawn at the widest point in the left ventricle (LV). Endocardial and epicardial wall movement was traced for three replicate measurements of three cardiac cycles. The same short axis B mode cine loops were then analyzed using VevoStrain. This method was repeated for three measurements (baseline, Frank-Starling, Sympathetic Response). Short axis Radial and Circumferential directions of movement were obtained. Strain was used as a measure of systolic function, while strain rate was used as contractility as it exhibits less load dependence.^21^ The reverse peak of strain rate was used as a measure of ventricular diastolic function.^21^ This method was repeated for three experimental manipulations (baseline, preload, and dobutamine).

### *In vivo* assessment of contractile reserve

After obtaining baseline echocardiography measurements, sterile saline (150 μL, Hospira, Lake Forest, IL) was injected intravenously (either through tail vein or retro orbital vein) and B mode images were immediately obtained to investigate Frank-Starling Law of the Heart (referred to as “preload” measurements). After recovery, mice were then administered the β1-AR agonist dobutamine (50 μL, 1 mg/mL, Hospira, Lake Forest, IL) intravenously (either tail or retro orbital vein). B-mode images were immediately obtained to investigate the sympathetic response. All images (Baseline, Preload, and Sympathetic Stimulation) were obtained in the same viewing location in the heart. Mice were allowed to recover before return to their cage. Cardiac reserve was measured by response of all cardiac parameters. Contractile reserve was measured through response in strain rate (radial and circumferential) with increased saline or dobutamine injection compared to baseline.

### Data exclusion

Data exclusion criteria were created before the experiment was performed: Mice must have analyzable images (entire left ventricle visible with no rib or sternum shadow) and demonstrate an increase in preload with saline injection. One (1) control mouse was excluded for preload data due to a lack of increase in stroke volume. Baseline and dobutamine values were still used. One (1) HFD mouse was excluded in preload data due to poor image acquisition with echocardiography. Baseline and dobutamine values were still used. One (1) aging mouse was excluded as a statistical outlier (Grubbs’ test, α= 0.05) for baseline, preload, and dobutamine. Individuals making these decisions were not blinded from experimental group.

### Blinding

Due to the nature of the mouse models, the two echocardiographers were not blinded during acquisition of data. Analyzers were blinded for group (i.e., control, aging, HFD, exercise-training) but not treatment (baseline, preload, dobutamine).

### Statistical tests

Baseline data was analyzed by student t-test against control mice for all groups. Frank-Starling and Sympathetic response data was analyzed via 2 way ANOVA. Post hoc Sidak’s comparison were used to compared between groups for ANOVA calculations. Data in supplementary tables are shown as mean ± SEM.

## RESULTS

### Assessing contractile reserve with STE in mice

Previous studies have established that exercise training results in the heart having an enhanced functional response to β-AR stimulation.^24^ We therefore used an HIIT protocol to exercise train mice; these mice were used as a positive control to establish a non-invasive, non-terminal procedure for assessing contractile reserve *in vivo* through echocardiography. At baseline, exercise-trained mice exhibited no differences in cardiovascular performance (measured as ejection fraction), heart performance (measured as stroke volume), systolic function (measured as circumferential and radial strain), diastolic function (measured as reverse circumferential and radial strain), and contractility (measured as circumferential and radial strain rate) compared to control mice (Supplementary Table 1). However, exercise mice did have increased LV diastolic posterior wall thickness and increased LV mass compared to control mice (Supplementary Table 1), consistent with previous observations of physiological hypertrophy in athletic hearts.^24^

Frank-Starling response was assessed by saline injection in both control and exercise trained mice. We observed an increase in preload measured as end diastolic volume (Figure 1A). The Frank-Starling mechanism is based on the premise that an increase in preload will cause a corresponding increase in contraction and therefore we next looked at stroke volume. Heart performance, measured as stroke volume, increased in control but not exercise-trained mice (Figure 1B). Consistent with strain and strain rate being virtually load-independent, we observed no change in measurements of systolic and diastolic function (Figure 1C and Supplementary Table 2). Interestingly, contractility (circumferential strain rate) decreased with increased preload with exercise (Figure 1C), but stroke volume was still maintained, consistent with Frank-Starling Law of the Heart being more dominant in the exercise heart.^27^ We speculate that the decrease in contractility is likely due to the fact that exercise mice are highly reliant upon the preload mechanism^28, 29^. Since we have artificially increased preload; the heart does not need to work as hard because of this to maintain overall performance and contractility was decreased.

**Figure 1:**
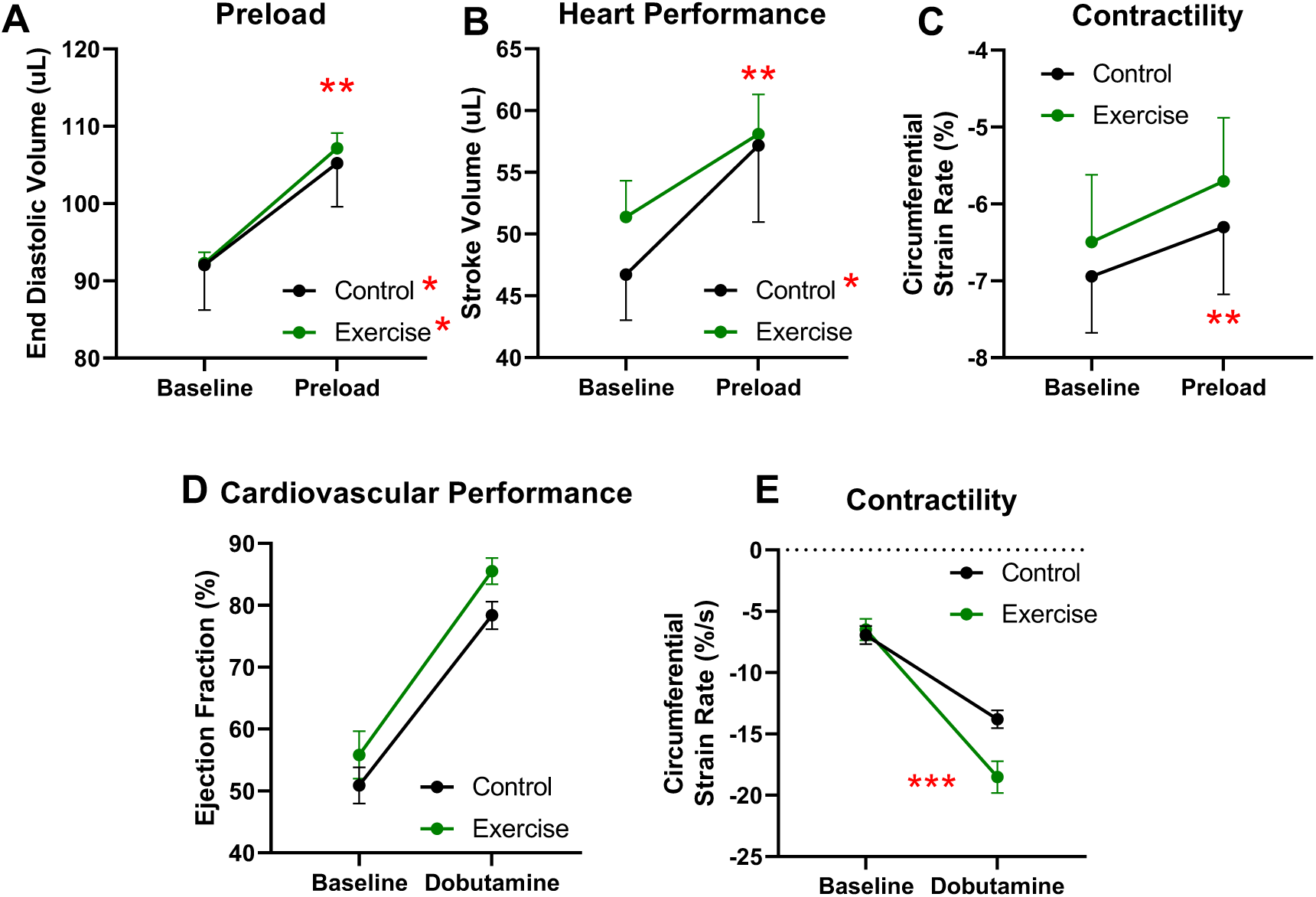
Echocardiography measurements of heart function at baseline, with increased preload, and with sympathetic stimulation for control and exercise-trained mice. Assessment of the Frank-Starling mechanism of the heart in control and exercise mice through (A) end diastolic volume, (B) stroke volume, and (C) contractility, measured as circumferential strain rate. Assessment of sympathetic response via intravenous dobutamine injection through (D) cardiovascular performance, measured as Ejection Fraction, and (E) contractility, measured as Circumferential Strain Rate. *p≤0.05 2-way ANOVA, Sidak’s comparison; **p≤0.05 2-way ANOVA, Preload injection vs. measurement; ***p≤0.05 2-way ANOVA, interaction. N=4-7 mice/group.

Consistent with previous studies^19^, in response to sympathetic stimulation with a β1-AR agonist, we observed an increase in cardiovascular performance (measured as ejection fraction) for both control and exercise-trained mice (Figure 1D). As exercise-training increases contractile reserve, we used STE to assess indices of systolic function, diastolic function, and contractility. Our data indicate that in response to dobutamine, exercise-trained mice demonstrate a significantly greater increase in contractility compared to control mice, measured as circumferential strain rate (Figure 1E), consistent with measurements performed using invasive technique,^18^ indicating an enhanced contractile reserve. Exercise-trained mice also demonstrated an increased chronotropic reserve evident through a greater increase in heart rate compared to control mice (Supplementary Table 3). Our data are consistent with exercise resulting in a greater cardiac reserve^30^. These data demonstrate that we have developed a non-terminal, non-invasive procedure for assessing cardiac reserve and assessing load-independent measurements of systolic function, diastolic function, and contractility in mice through STE.

### Poor systolic function, diastolic function, and contractility in very old mice

At baseline, very old mice did not have reduced cardiovascular performance (defined as <40% ejection fraction) but did display a trend towards a reduced ejection fraction compared to younger mice (Figure 2A). Additionally, stroke volume did not differ between aging and young mice (Supplementary Table 1). However, we did observe increased LV mass and diastolic posterior wall thickness indicating that these old mice have hypertrophic hearts (Figures 2B, 2C). We next examined systolic function, diastolic function, and contractility by performing the first ever studies using STE on mice of this age. Our data show that mice with advanced age have decreased systolic function and load-independent contractility compared to control mice measured via both circumferential parameters (Figure 2D-F) and radial parameters (Supplementary Table 1). There was also a trend that diastolic function (measured via reverse circumferential strain) and radial strain (Supplementary Table 1) was blunted. We believe the ejection fraction and stroke volume in these aging mice are maintained due to the remodeling of the heart. Thus, we further investigated the hypertrophy by examining the concentricity (LVPW;d/EDD) and found that mice with advanced age demonstrated increased concentric hypertrophy, consistent with previous studies.^31^ While global parameters of cardiovascular and heart function are maintained, STE revealed that systolic function, diastolic function, and contractility are all diminished with aging.

**Figure 2:**
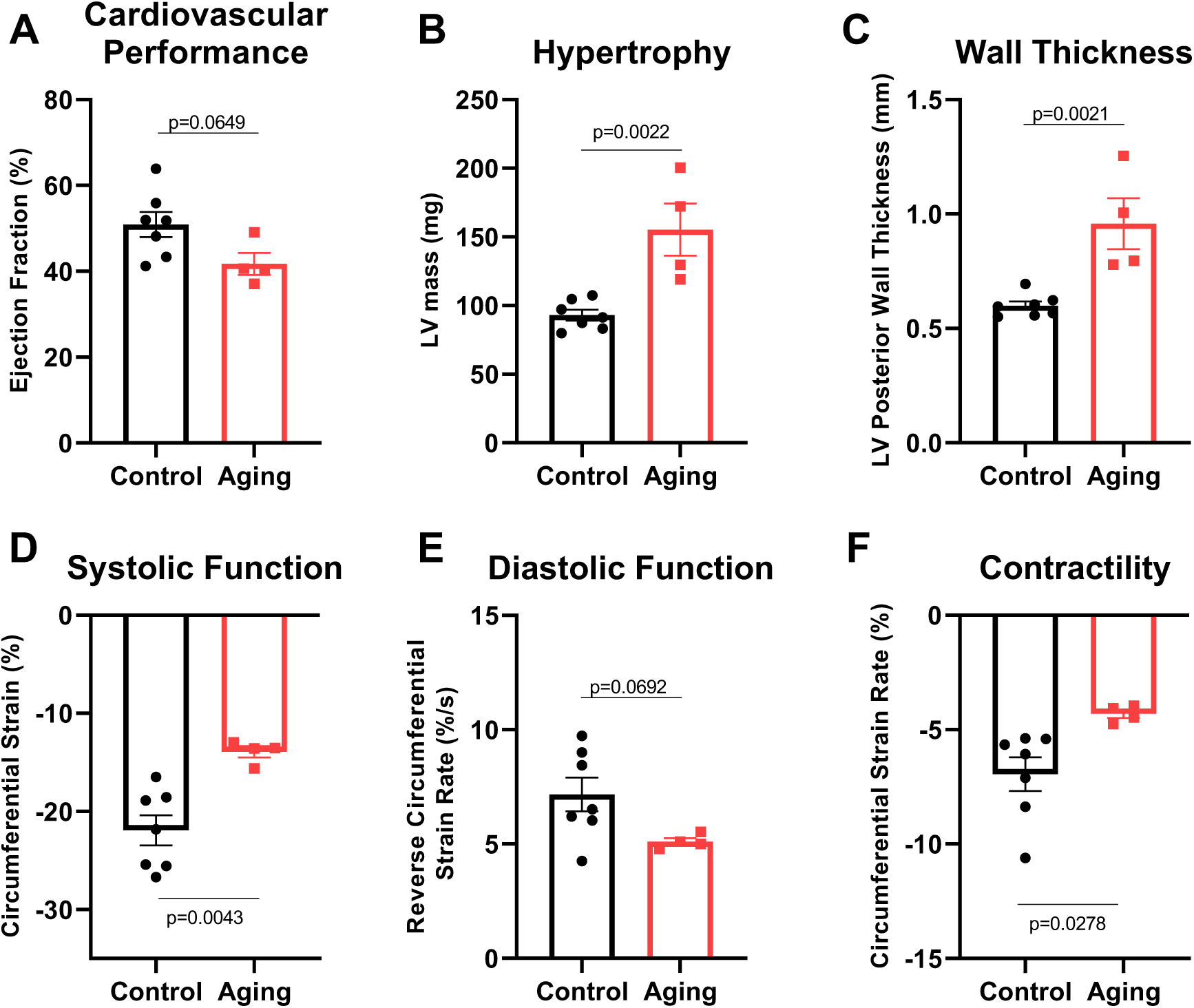
Echocardiography measurements of heart function at baseline for aged (24 month old) mouse model compared to control mice. (A) Cardiovascular performance measured as Ejection Fraction, (B) Hypertrophy measured as Left Ventricular (LV) mass, (C) Wall Thickness measured as LV posterior wall thickness, (D) Systolic function measured as circumferential strain, (E) Diastolic function measured as Reverse circumferential strain rate, and (F) Contractility measured as circumferential strain rate.. Statistical tests are student t-tests. N=4-7 mice/group.

### Aging mice exhibit a blunted contractile reserve

In contrast to both young control and exercise-trained mice, aging mice had no significant increase in either preload and stroke volume (Figure 3A, 3B) with saline injection. However, aging mice did exhibit an increase in cardiovascular performance (ejection fraction) in response to increased preload (Figure 3C). Aging mice had no change in strain or strain rates after increasing preload, indicating that these are still load independent parameters even in aging models. Systolic function, diastolic function, and contractility were still all blunted in aged mice compared to young mice with increasing preload (Figures 3D-F). Thus, mice with advanced age have a blunted Frank-Starling Law of the Heart.

**Figure 3:**
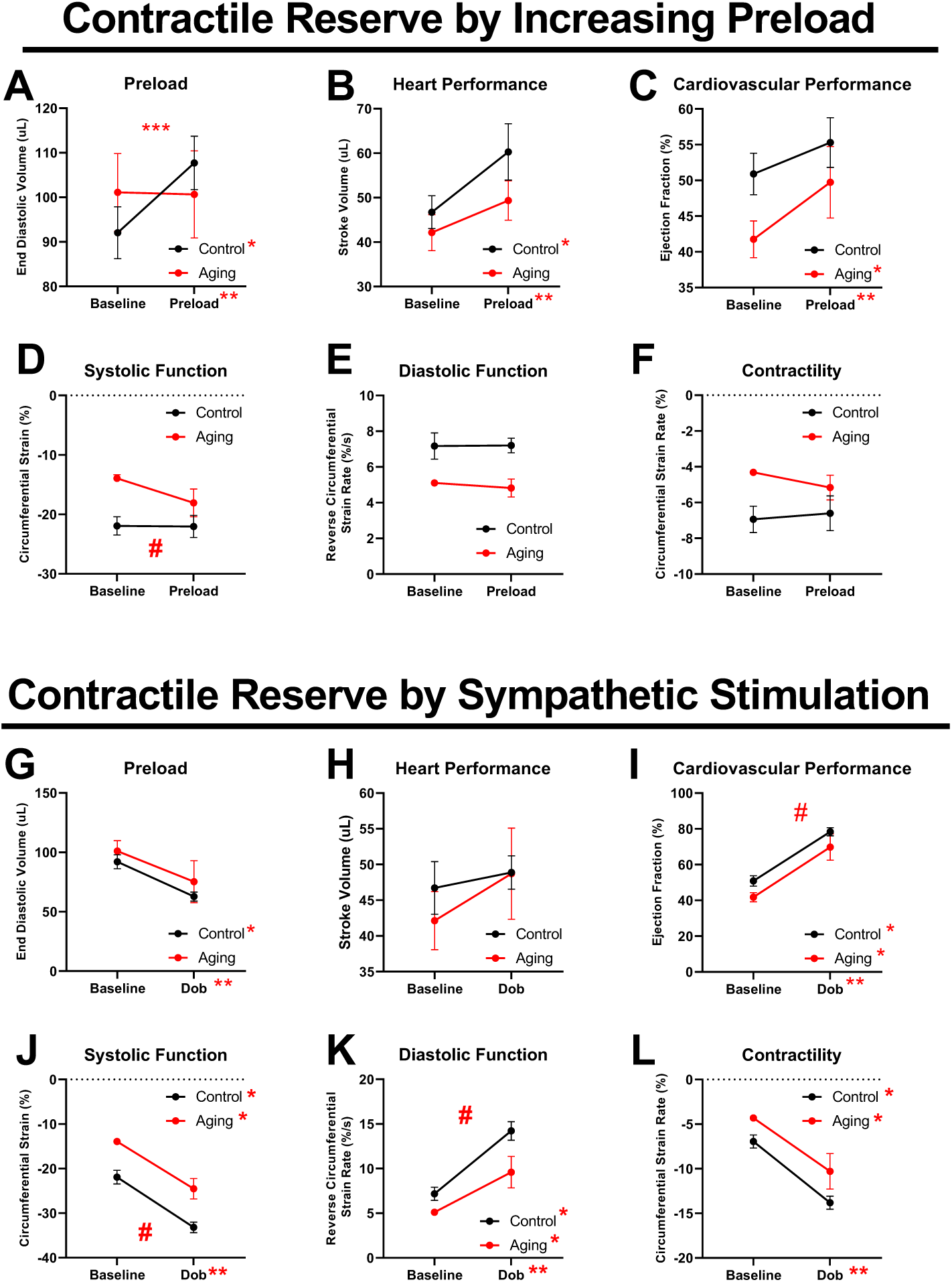
Echocardiography measurements of heart function compared to baseline for control and aging mice. Assessment of the Frank-Starling mechanism of the heart through (A) Preload measured as end diastolic volume, (B) heart performance measured by stroke volume, (C) Cardiovascular performance measured as Ejection Fraction, (D) Systolic function measured as circumferential strain, (E) Diastolic function measured as Reverse circumferential strain rate, and (F) Contractility measured as circumferential strain rate. Corresponding measurements were also performed in response to dobutamine (G-L). *p≤0.05 2-way ANOVA, Sidak’s comparison; **p≤0.05 2-way ANOVA, injection vs. measurement; ***p≤0.05 2-way ANOVA, interaction; # p≤0.05 2-way ANOVA, Control v. Aging. N=4-7 mice/group. Dob=dobutamine.

Dobutamine injection resulted in a significant decrease in end diastolic volume in both control and aging mice. This may be due to the increase in heart rate, leading to decreased time in diastole and less time for ventricular filling (Figure 3G). Even though preload was decreased, heart performance (measured as stroke volume) was unaltered compared to baseline in both control and aging mice (Figure 3H). Overall, this contributed to a blunted cardiovascular performance with sympathetic stimulation in aging mice, measured as ejection fraction (Figure 3I). Further, sympathetic stimulation did increase systolic function, diastolic function and contractility in aging mice compared to baseline. However, this response was blunted compared to young control mice (circumferential- Figure 3J-L; radial- Supplementary Table 5). Interestingly, diastolic function was not accelerated in aging mice with sympathetic stimulation (circumferential- Figure 3K; radial- Supplementary Table 5). These are the first data using STE to demonstrate that contractile reserve is blunted in mice with advanced age.

### Poor systolic function, diastolic function, and contractility in obese mice

Cardiovascular (measured as ejection fraction) and heart (measured as stroke volume) performance in HFD mice were not different from control mice (Figure 4A and Supplementary Table 1). However, HFD mice demonstrated increased LV mass and wall thicknesses, consistent with hypertrophy observed in the HFD mouse model (Figure 4B, 4C).^18^ Our data also revealed that HFD mice demonstrated decreased circumferential systolic function, diastolic function, and contractility (Figure 4D-F). Radial measurements were decreased compared to control as well (Supplementary Table 1). Since global cardiovascular parameters were maintained in obesity, we believe that this is due to the remodeling of the heart. HFD mice demonstrated hypertrophy through increased wall thickness and LV mass compared to control, yet did not have significant concentricity (Supplementary Table 1). This is consistent with previous reports demonstrating that obesity results in eccentric hypertrophy.^32^ While global parameters of cardiovascular and heart function are maintained, STE revealed that systolic function, diastolic function, and contractility are all diminished with obesity.

**Figure 4:**
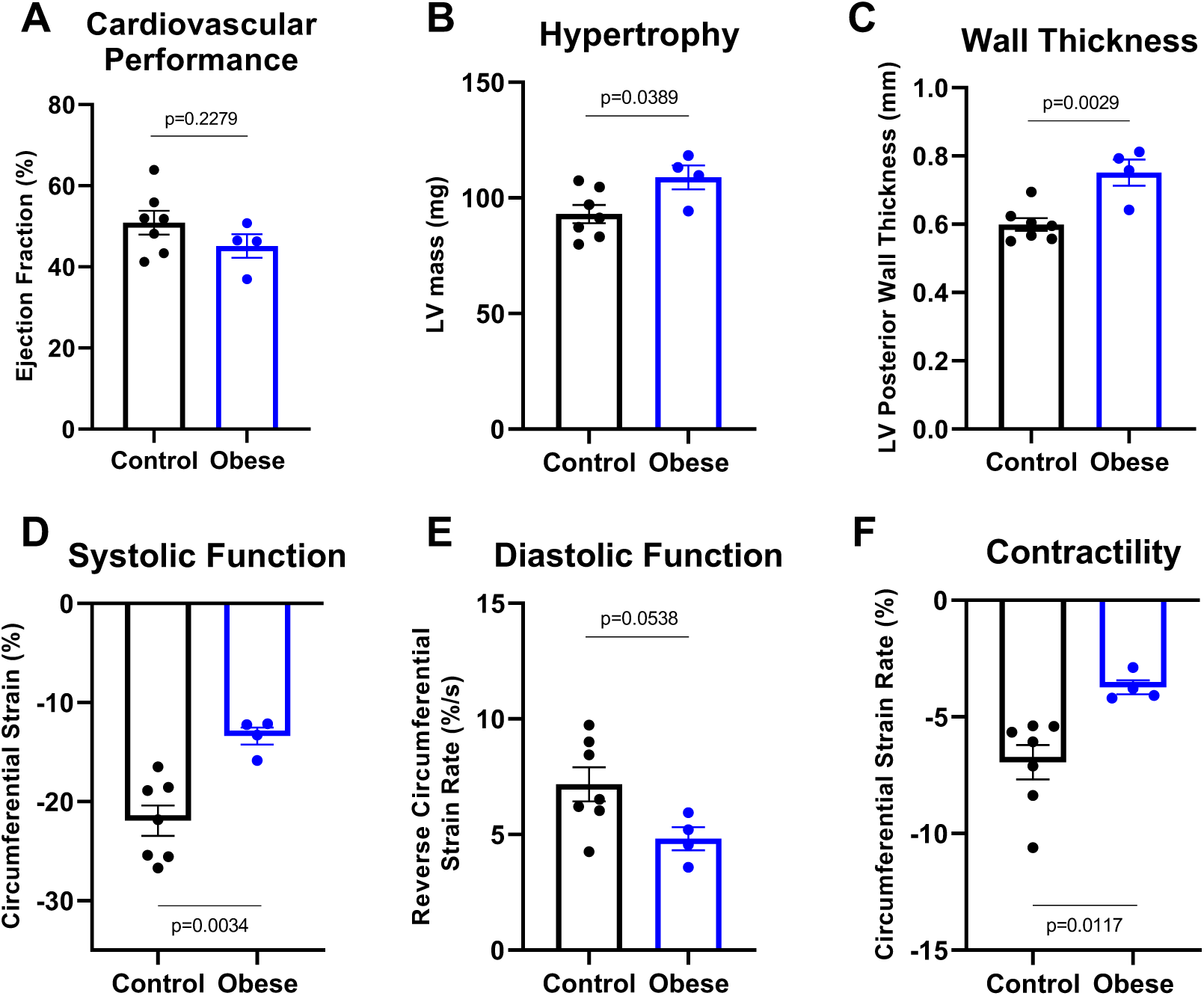
Echocardiography measurements of heart function at baseline for obese (high fat diet-induced) mouse model compared to control mice. (A) Cardiovascular performance measured as Ejection Fraction, (B) Hypertrophy measured as Left Ventricular (LV) mass, (C) Wall Thickness measured as LV posterior wall thickness, (D) Systolic function measured as circumferential strain, (E) Diastolic function measured as Reverse circumferential strain rate, and (F) Contractility measured as circumferential strain rate. Statistical tests are student t-tests. N=4-7 mice/group.

### Obese mice exhibit a blunted contractile reserve

Similar to control mice, HFD had a significant increase in both preload (end diastolic volume) and stroke volume in response to saline injection (Figure 5A, 5B). As a result of this, HFD mice had increased cardiovascular performance measured as ejection fraction (Figure 5C). Unlike aging mice, HFD mice exhibited increased systolic function, diastolic function, and contractility with increased preload; Although, this response was still blunted compared to control mice with increasing preload (radial – Supplementary Table 3, circumferential- Figure 5D-F). Interestingly, in contrast to control and exercise-trained mice, HFD exhibited increases in load-independent measurements. The observed increase in systolic function in the radial direction after increased preload (see Supplementary Table 3) may be the reason why Frank-Starling Law of the Heart is maintained in obesity.

**Figure 5:**
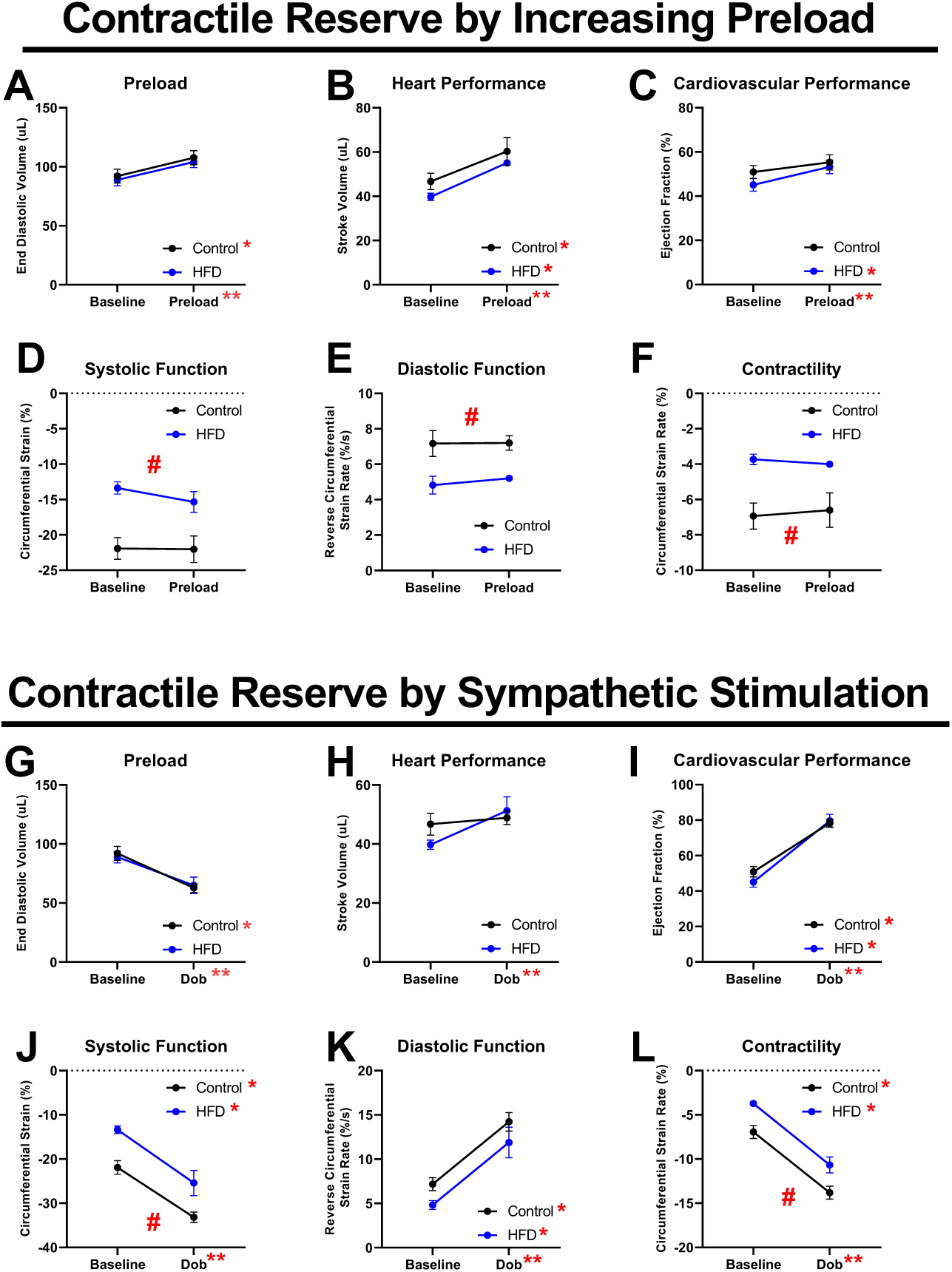
Echocardiography measurements of heart function compared to baseline for control and obese (HFD) mice. Assessment of the Frank-Starling mechanism of the heart through (A) Preload measured as end diastolic volume, (B) heart performance measured by stroke volume, (C) Cardiovascular performance measured as Ejection Fraction, (D) Systolic function measured as circumferential strain, (E) Diastolic function measured as Reverse circumferential strain rate, and (F) Contractility measured as circumferential strain rate. Corresponding measurements were also performed in response to dobutamine (G-L). *p≤0.05 2-way ANOVA, Sidak’s comparison; **p≤0.05 2- way ANOVA, injection vs. measurement; ***p≤0.05 2-way ANOVA, interaction; # p≤0.05 2-way ANOVA, Control v. HFD. N=3-7 mice/group based on treatment. Dob=dobutamine.

In response to sympathetic stimulation, there was a trend that HFD mice had a decrease in end diastolic volume, similar to control and aging mice (Figure 5G). Dobutamine also resulted in an increase in cardiovascular (ejection fraction) and heart (stroke volume) performance (Figure 5H, 5I). While sympathetic activation in HFD mice did increase systolic function, diastolic function, and contractility compared to baseline, this response was significantly blunted compared to control mice (circumferential parameters- Figure 5J-L, radial parameters- Supplemental Table 5). These are the first data using STE to demonstrate that contractile reserve is blunted in HFD mice and different than aging mice.

### Obese and Aging mice have different contractile reserves

Since we observed differences in contractile reserve between aging and HFD mice, we further investigated the characteristics of myocardial contractile function by examining wall displacement.^33^ Our data show that in response to increasing preload, aging mice had both greater radial endocardial and greater epicardial wall displacement (in relation to strain, Figure 6A-C), while HFD mice had significantly greater circumferential endocardial and epicardial wall displacement (Figure 6D-F). There was no difference between aging and HFD in radial endocardial and epicardial wall displacement in response to sympathetic stimulation (Figure 6G-I). However, HFD mice had significantly greater circumferential endocardial and epicardial wall displacement after sympathetic stimulation compared to aging mice (Figure 6J-L). We believe these directional differences in wall displacement contribute to the model-specific response to increased preload and sympathetic stimulation.

**Figure 6:**
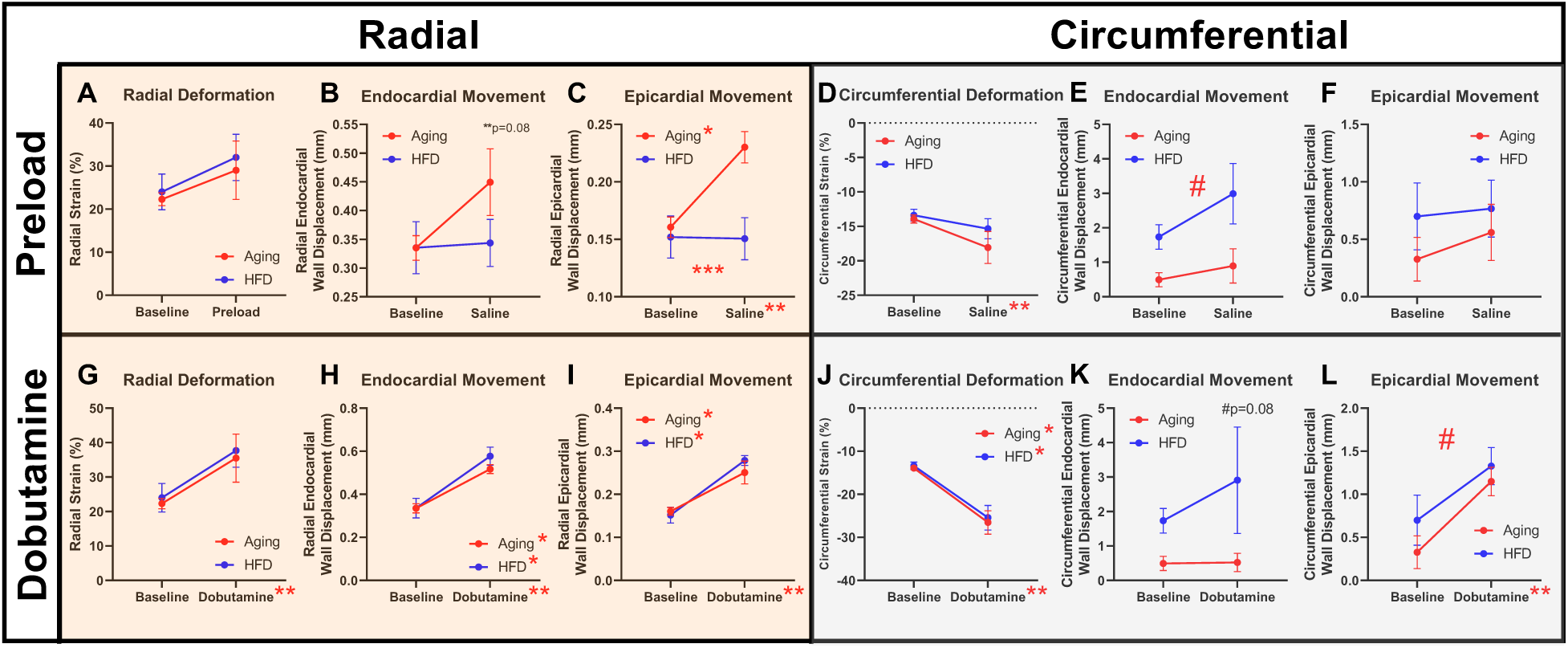
**Strain and wall displacement for aging and obese mice**. Assessment of the Frank-Starling mechanism of the heart through in circumferential and radial directions by (A) Radial deformation measured as radial strain, (B) Endocardial movement measured as radial epicardial wall displacement, (C) Epicardial movement measured as radial epicardial wall displacement, (D) circumferential deformation measured as circumferential strain, (E) Endocardial movement measured as circumferential endocardial wall displacement, and (F) Epicardial movement measured as circumferential epicardial wall displacement. Corresponding measurements were also performed in response to dobutamine (G-L). *p≤0.05 2-way ANOVA, Sidak’s comparison; **p≤0.05 2- way ANOVA, injection vs. measurement; ***p≤0.05 2-way ANOVA, interaction; # p≤0.05 2-way ANOVA, Aging v. HFD. N=3-4 mice/group based on treatment.

## DISCUSSION

Contractile reserve is an essential attribute of the heart and is necessary to increase function in response to heightened metabolic demand. Unfortunately, in the context of heart disease, contractile reserve is blunted and emerges as a key diagnostic indicator of disease incidence and progression.^34–36^ Contractile reserve has not been fully characterized in murine models of aging and obesity. Traditional methods of assessing contractile reserve in animal models, such as intra left ventricular catheterization, pose limitations due to their terminal nature and output of load-and heart rate-dependent indices which complicates data interpretation.^21^ Given the shift in human population demographics, particularly in the rise in elderly and obese individuals^2, 5^, a non-terminal, load-independent measure of contractile reserve in mouse models becomes imperative. Thus, we utilized STE to investigate the heart function of murine models of aging and obesity, expanding previous characterizations of these models by including assessment of contractility, diastolic function, and contractile reserve.^37, 38^

In this study, we mark the first investigation of comprehensive STE in murine models of aging and obesity. Consistent with previous studies, global parameters of cardiovascular and heart performance (ejection fraction and stroke volume) were not reduced in aging and obese mice. However, focused analyses of baseline systolic function, diastolic function, and contractility revealed blunted function in both cohorts compared to control mice (Figures 2 and 4, Supplementary Table 1). Notably, when comparing the contractile reserve, we observed differences between aging mice and obese mice (Figure 3, 5 and 6). Despite strain and strain rate exhibiting load independence in control and exercise trained mice (serving as a model of increased cardiac function), obese mice showed increased systolic and diastolic function and contractility compared to baseline in response to increased load (Figure 3), delineating distinct mechanisms underlying contractile reserve in these disease models. Interestingly, this suggests that contractile reserve is being accessed in obese mice at baseline and, upon increased demand via increased preload, the heart must employ other mechanisms to respond to increased preload as compared to control and exercise mice. This may explain why obese mice are more reliant on Frank-Starling to increase their heart performance^39^.

With this dichotomy in response to increasing contractile reserve, we further investigated wall displacement. Both wall displacement and strain are measurements used to describe cardiac function, yet they represent different aspects of LV mechanics.^33^ While displacement refers to the absolute movement of the myocardial wall during the cardiac cycle, strain quantifies tissue deformation relative to its initial length, thus offering insights into contractile function of the heart. We found that obese mice had greater circumferential wall displacement (in relation to strain) compared to aging mice, while aging mice had enhanced radial endocardial and epicardial wall displacement after increasing preload compared to obese mice (Figure 6).

Our findings revealed distinct patterns of wall displacement between aging and obese mice, suggesting differing mechanisms of remodeling underlying these disparities, potentially driven by fibrosis^40, 41^. Fibrosis of the heart is an essential compensatory mechanism in disease (e.g., HFpEF) ^42^ and while fibrosis is found in both aging and obese animals and humans^43, 44^, it has been shown that location of fibrosis within the heart differentially impacts mechanical deformation.^42^ The influence of fibrosis also extends to contractile reserve, as evidenced by studies demonstrating a strong correlation between the extent of histological disruption and contractile reserve^45^. Another potential explanation for our observations may be due to cardiac fiber orientation. Animal models of non-ischemic heart disease demonstrate changes in myocardial fiber orientation and remodeling between healthy and diseased hearts.^46, 47^ Therefore, variation in the endocardial and epicardial wall movement may be due to changes in localized fiber orientation based on disease etiology. Our study demonstrates functional disparities between contractile reserve response in aging and obesity that may be attributed to variations in remodeling, such as hypertrophy, cardiac fiber orientation, and fibrosis.

A limitation of this study is the sample sizes of experimental groups. However, given the statistically-distinct phenotypes observed, we do not anticipate that our conclusions would be different with a larger sample size. We demonstrate distinct functional phenotypes between aged and obese mice that do not have impaired ejection fraction, yet demonstrate impaired heart function and blunted contractile reserves. While we did not investigate the tissue of these mice for markers of HFpEF, we believe these differences contribute to the distinct HFpEF phenotypes observed with aging and obesity.^48, 49^

## CONCLUSION

In this study, we investigated the contractile reserve and LV mechanics in murine models of aging and obesity using STE. We developed a novel method to non-terminally and non-invasively assess contractile reserve *in vivo*. We found that, while global cardiac function appeared normal in both aging and obese mice, STE revealed attenuated systolic function, diastolic function, and contractility in both aging and obese mice. Interestingly, unlike aging mice, obese mice demonstrated increased cardiovascular and heart performance with increased preload. Moreover, aged and obese mice exhibited different patterns in wall displacement. These findings provide insights into the distinct mechanisms underlying contractile reserve and LV mechanics in aging and obesity-induced cardiac dysfunction, potentially elucidating the emergence of different heart failure phenotypes in these patient populations. The variations we observe in wall displacement and cardiac reserve between aging and obesity could potentially contribute to the characterization of these distinct phenotypes.

## Acknowledgements

### Sources of Funding

This work was supported by T32 HL134616 (S.L. Sturgill) and R01 AG060542 (M.T. Ziolo).

### Disclosures

None

### Supplemental Materials

Tables S1-S7

**References**

**Tables**

## References

1. Woodruff RC, Tong X, Khan SS, Shah NS, Jackson SL, Loustalot F and Vaughan AS. Trends in Cardiovascular Disease Mortality Rates and Excess Deaths, 2010-2022. American journal of preventive medicine. 2024;66:582-589.

2. Anderson LA, Goodman RA, Holtzman D, Posner SF and Northridge ME. Aging in the United States: opportunities and challenges for public health. American journal of public health. 2012;102:393–5.

3. Yazdanyar A and Newman AB. The burden of cardiovascular disease in the elderly: morbidity, mortality, and costs. Clinics in geriatric medicine. 2009;25:563–77, vii.

4. Powell-Wiley TM, Poirier P, Burke LE, Després J-P, Gordon-Larsen P, Lavie CJ, Lear SA, Ndumele CE, Neeland IJ, Sanders P and St-Onge M-P. Obesity and Cardiovascular Disease: A Scientific Statement From the American Heart Association. Circulation. 2021;143:e984–e1010.

5. Ward ZJ, Bleich SN, Cradock AL, Barrett JL, Giles CM, Flax C, Long MW and Gortmaker SL. Projected U.S. State-Level Prevalence of Adult Obesity and Severe Obesity. The New England journal of medicine. 2019;381:2440–2450.

6. Borlaug BA, Jensen MD, Kitzman DW, Lam CSP, Obokata M and Rider OJ. Obesity and heart failure with preserved ejection fraction: new insights and pathophysiological targets. Cardiovascular research. 2023;118:3434–3450.

7. Upadhya B, Taffet GE, Cheng CP and Kitzman DW. Heart failure with preserved ejection fraction in the elderly: scope of the problem. J Mol Cell Cardiol. 2015;83:73–87.

8. Dădârlat-Pop A, Sitar-Tăut A, Zdrenghea D, Caloian B, Tomoaia R, Pop D and Buzoianu A. Profile of Obesity and Comorbidities in Elderly Patients with Heart Failure. Clinical interventions in aging. 2020;15:547–556.

9. Ohte N. Heart Failure with Preserved Ejection Fraction Is a Still Big Unmet Need in Cardiology. Journal of clinical medicine. 2021;10.

10. Xue J, Zhao F, Wang Y, Gu J, Gao J, Wang X and Zhou H. Integrative Cardiac Reserve. Integrative Medicine International. 2015;1:162–169.

11. Francis GS and Desai MY. Contractile reserve: are we beginning to understand it? JACC Cardiovascular imaging. 2008;1:727–8.

12. Di Lisi D, Ciampi Q, Madaudo C, Manno G, Macaione F, Novo S and Novo G. Contractile Reserve in Heart Failure with Preserved Ejection Fraction. Journal of cardiovascular development and disease. 2022;9.

13. Borlaug BA, Olson TP, Lam CS, Flood KS, Lerman A, Johnson BD and Redfield MM. Global cardiovascular reserve dysfunction in heart failure with preserved ejection fraction. Journal of the American College of Cardiology. 2010;56:845–54.

14. Tan LB and Littler WA. Measurement of cardiac reserve in cardiogenic shock: implications for prognosis and management. Br Heart J. 1990;64:121–8.

15. Foulkes S, Claessen G, Howden EJ, Daly RM, Fraser SF and La Gerche A. The Utility of Cardiac Reserve for the Early Detection of Cancer Treatment-Related Cardiac Dysfunction: A Comprehensive Overview. Frontiers in cardiovascular medicine. 2020;7:32.

16. Moss RL and Fitzsimons DP. Frank-Starling relationship: long on importance, short on mechanism. Circulation research. 2002;90:11–3.

17. Bers DM and Ziolo MT. When is cAMP not cAMP? Effects of compartmentalization. Circulation research. 2001;89:373–5.

18. Calligaris SD, Lecanda M, Solis F, Ezquer M, Gutiérrez J, Brandan E, Leiva A, Sobrevia L and Conget P. Mice long-term high-fat diet feeding recapitulates human cardiovascular alterations: an animal model to study the early phases of diabetic cardiomyopathy. PloS one. 2013;8:e60931.

19. Haggerty CM, Mattingly AC, Kramer SP, Binkley CM, Jing L, Suever JD, Powell DK, Charnigo RJ, Epstein FH and Fornwalt BK. Left ventricular mechanical dysfunction in diet-induced obese mice is exacerbated during inotropic stress: a cine DENSE cardiovascular magnetic resonance study. Journal of cardiovascular magnetic resonance : official journal of the Society for Cardiovascular Magnetic Resonance. 2015;17:75.

20. Kovács A, Oláh A, Lux Á, Mátyás C, Németh BT, Kellermayer D, Ruppert M, Török M, Szabó L, Meltzer A, Assabiny A, Birtalan E, Merkely B and Radovits T. Strain and strain rate by speckle-tracking echocardiography correlate with pressure-volume loop-derived contractility indices in a rat model of athlete’s heart. American journal of physiology Heart and circulatory physiology. 2015;308:H743–8.

21. Sturgill SL, Shettigar V and Ziolo MT. Antiquated ejection fraction: Basic research applications for speckle tracking echocardiography. Frontiers in physiology. 2022;13:969314.

22. Frederiksen PH, Linde L, Gregers E, Udesen NLJ, Helgestad OK, Banke A, Dahl JS, Povlsen AL, Jensen LO, Larsen JP, Lassen J, Schmidt H, Ravn HB and Moller JE. Association between speckle tracking echocardiography and pressure-volume loops during cardiogenic shock development. Open Heart. 2024;11:e002512.

23. Roof SR, Ho HT, Little SC, Ostler JE, Brundage EA, Periasamy M, Villamena FA, Gyorke S, Biesiadecki BJ, Heymes C, Houser SR, Davis JP and Ziolo MT. Obligatory role of neuronal nitric oxide synthase in the heart’s antioxidant adaptation with exercise. Journal of molecular and cellular cardiology. 2015;81:54–61.

24. Roof SR, Tang L, Ostler JE, Periasamy M, Gyorke S, Billman GE and Ziolo MT. Neuronal nitric oxide synthase is indispensable for the cardiac adaptive effects of exercise. Basic Res Cardiol. 2013;108:332.

25. Peres Valgas da Silva C, Shettigar VK, Baer LA, Abay E, Madaris KL, Mehling MR, Hernandez-Saavedra D, Pinckard KM, Seculov NP, Ziolo MT and Stanford KI. Brown adipose tissue prevents glucose intolerance and cardiac remodeling in high-fat-fed mice after a mild myocardial infarction. International journal of obesity (2005). 2022;46:350-358.

26. Pinckard KM, Shettigar VK, Wright KR, Abay E, Baer LA, Vidal P, Dewal RS, Das D, Duarte- Sanmiguel S, Hernández-Saavedra D, Arts PJ, Lehnig AC, Bussberg V, Narain NR, Kiebish MA, Yi F, Sparks LM, Goodpaster BH, Smith SR, Pratley RE, Lewandowski ED, Raman SV, Wold LE, Gallego-Perez D, Coen PM, Ziolo MT and Stanford KI. A Novel Endocrine Role for the BAT-Released Lipokine 12,13-diHOME to Mediate Cardiac Function. Circulation. 2021;143:145–159.

27. La Gerche A and Gewillig M. What Limits Cardiac Performance during Exercise in Normal Subjects and in Healthy Fontan Patients? International journal of pediatrics. 2010;2010.

28. Bevilacqua M, Savonitto S, Bosisio E, Chebat E, Bertora PL, Sardina M and Norbiato G. Role of the Frank-Starling mechanism in maintaining cardiac output during increasing levels of treadmill exercise in beta-blocked normal men. Am J Cardiol. 1989;63:853–7.

29. Horwitz LD, Atkins JM and Leshin SJ. Role of the Frank-Starling Mechanism In Exercise. Circulation research. 1972;31:868–875.

30. Le TT, Bryant JA, Ting AE, Ho PY, Su B, Teo RC, Gan JS, Chung YC, O’Regan DP, Cook SA and Chin CW. Assessing exercise cardiac reserve using real-time cardiovascular magnetic resonance. Journal of cardiovascular magnetic resonance : official journal of the Society for Cardiovascular Magnetic Resonance. 2017;19:7.

31. Ganau A, Saba PS, Roman MJ, de Simone G, Realdi G and Devereux RB. Ageing induces left ventricular concentric remodelling in normotensive subjects. Journal of hypertension. 1995;13:1818–22.

32. Cuspidi C, Rescaldani M, Sala C and Grassi G. Left-ventricular hypertrophy and obesity: a systematic review and meta-analysis of echocardiographic studies. Journal of hypertension. 2014;32:16–25.

33. Bijnens B, Claus P, Weidemann F, Strotmann J and Sutherland GR. Investigating cardiac function using motion and deformation analysis in the setting of coronary artery disease. Circulation. 2007;116:2453–64.

34. Kobayashi M, Izawa H, Cheng XW, Asano H, Hirashiki A, Unno K, Ohshima S, Yamada T, Murase Y, Kato TS, Obata K, Noda A, Nishizawa T, Isobe S, Nagata K, Matsubara T, Murohara T and Yokota M. Dobutamine stress testing as a diagnostic tool for evaluation of myocardial contractile reserve in asymptomatic or mildly symptomatic patients with dilated cardiomyopathy. JACC Cardiovascular imaging. 2008;1:718–26.

35. Thein PM, Mirzaee S, Cameron JD and Nasis A. Left ventricular contractile reserve as a determinant of adverse clinical outcomes: a systematic review. Internal medicine journal. 2022;52:186–197.

36. Waddingham PH, Bhattacharyya S, Zalen JV and Lloyd G. Contractile reserve as a predictor of prognosis in patients with non-ischaemic systolic heart failure and dilated cardiomyopathy: a systematic review and meta-analysis. Echo research and practice. 2018;5:1–9.

37. Zhang X, Kong S, Wu M, Niu Y, Wang K, Zhu H and Yuan J. Impact high fat diet on myocardial strain in mice by 2D speckle tracking imaging. Obesity research & clinical practice. 2021;15:133–137.

38. de Lucia C, Wallner M, Eaton DM, Zhao H, Houser SR and Koch WJ. Echocardiographic Strain Analysis for the Early Detection of Left Ventricular Systolic/Diastolic Dysfunction and Dyssynchrony in a Mouse Model of Physiological Aging. *The journals of gerontology Series A*, Biological sciences and medical sciences. 2019;74:455–461.

39. Vasan RS. Cardiac function and obesity. Heart (British Cardiac Society). 2003;89:1127–9.

40. Otasević P, Popović ZB, Vasiljević JD, Vidaković R, Pratali L, Vlahović A and Nesković AN. Relation of myocardial histomorphometric features and left ventricular contractile reserve assessed by high-dose dobutamine stress echocardiography in patients with idiopathic dilated cardiomyopathy. European journal of heart failure. 2005;7:49–56.

41. Collins J, Sommerville C, Magrath P, Spottiswoode B, Freed BH, Benzuly KH, Gordon R, Vidula H, Lee DC, Yancy C, Carr J and Markl M. Extracellular volume fraction is more closely associated with altered regional left ventricular velocities than left ventricular ejection fraction in nonischemic cardiomyopathy. Circulation Cardiovascular imaging. 2015;8.

42. Sweeney M, Corden B and Cook SA. Targeting cardiac fibrosis in heart failure with preserved ejection fraction: mirage or miracle? EMBO molecular medicine. 2020;12:e10865.

43. Biernacka A and Frangogiannis NG. Aging and Cardiac Fibrosis. Aging and disease. 2011;2:158–173.

44. Kruszewska J, Cudnoch-Jedrzejewska A and Czarzasta K. Remodeling and Fibrosis of the Cardiac Muscle in the Course of Obesity-Pathogenesis and Involvement of the Extracellular Matrix. International journal of molecular sciences. 2022;23.

45. Nagueh SF, Mikati I, Weilbaecher D, Reardon MJ, Al-Zaghrini GJ, Cacela D, He ZX, Letsou G, Noon G, Howell JF, Espada R, Verani MS and Zoghbi WA. Relation of the contractile reserve of hibernating myocardium to myocardial structure in humans. Circulation. 1999;100:490–6.

46. Wong J and Kuhl E. Generating fibre orientation maps in human heart models using Poisson interpolation. Computer methods in biomechanics and biomedical engineering. 2014;17:1217–26.

47. Schmitt B, Fedarava K, Falkenberg J, Rothaus K, Bodhey NK, Reischauer C, Kozerke S, Schnackenburg B, Westermann D, Lunkenheimer PP, Anderson RH, Berger F and Kuehne T. Three-dimensional alignment of the aggregated myocytes in the normal and hypertrophic murine heart. Journal of applied physiology (Bethesda, Md : 1985). 2009;107:921-7.

48. Obokata M, Reddy YNV, Pislaru SV, Melenovsky V and Borlaug BA. Evidence Supporting the Existence of a Distinct Obese Phenotype of Heart Failure With Preserved Ejection Fraction. Circulation. 2017;136:6–19.

49. Kitzman DW and Shah SJ. The HFpEF Obesity Phenotype: The Elephant in the Room. Journal of the American College of Cardiology. 2016;68:200–3.

