## Supplemental Data for "Old mice have a functionally distinct contractile reserve from obese mice"

|  | **Supplemental Table 1: Baseline Echocardiography** | | | |
| --- | --- | --- | --- | --- |
|  | **Parameter** | **Group** | **Mean SEM** | **Significance (t-test)** |
|  | Radial Strain (%) | Control | 41.1752 ± 2.6792 | - |
|  |  | Aging | 22.2985 ± 1.4801 | * |
|  |  | HFD | 24.0017 ± 4.1421 | * |
|  |  | Exercise | 37.4612 ± 2.3462 | NS |
|  | Radial Strain Rate (%/s) | Control | 8.2263 ± 0.3651 | - |
|  |  | Aging | 5.1116 ± 0.1581 | * |
|  |  | HFD | 4.8593 ± 0.5619 | * |
|  |  | Exercise | 7.9205 ± 0.2810 | NS |
|  | Reverse Peak Radial Strain Rate (%/s) | Control | -9.5950 ± 0.6965 | - |
|  |  | Aging | -7.2435 ± 0.8150 | NS (0.06) |
|  |  | HFD | -5.8448 ± 0.7917 | * |
|  |  | Exercise | -8.6798 ± 0.5834 | NS |
|  | Radial Endocardial Displacement (mm) | Control | 0.4247 ± 0.0236 | - |
|  |  | Aging | 0.3351 ± 0.0214 | * |
|  |  | HFD | 0.3354 ± 0.0453 | NS (0.08) |
|  |  | Exercise | 0.4271 ± 0.0135 | NS |
|  | Radial Epicardial Displacement (mm) | Control | 0.1788 ± 0.0214 | - |
|  |  | Aging | 0.161 ± 0.009 | NS |
|  |  | HFD | 0.1520 ± 0.0183 | NS |
|  |  | Exercise | 0.1751 ± 0.0194 | NS |
|  | Circumferential Strain (%) | Control | -21.9251 ± 1.5297 | - |
|  |  | Aging | -13.9298 ± 0.5841 | * |
|  |  | HFD | -13.3811 ± 0.8653 | * |
|  |  | Exercise | -18.7890 ± 1.2244 | NS |
|  | Circumferential Strain Rate (%/s) | Control | -6.9416 ± 0.7348 | - |
|  |  | Aging | -4.3120 ± 0.1854 | * |
|  |  | HFD | -3.7315 ± 0.2978 | * |
|  |  | Exercise | -6.4961 ± 0.8760 | NS |
|  | Reverse Circumferential Strain Rate (%/s) | Control | 7.1711 ± 0.7357 | - |
|  |  | Aging | 5.1029 ± 0.1568 | NS (0.06) |
|  |  | HFD | 4.8215 ± 0.5000 | NS (0.05) |
|  |  | Exercise | 5.9698 ± 0.4722 | NS |
|  | Circumferential Endocardial Displacement (mm) | Control | 1.803 ± 0.482 | - |
|  |  | Aging | 0.4937 ± 0.2037 | NS (0.08) |
|  |  | HFD | 1.734 ± 0.356 | NS |
|  |  | Exercise | 3.071 ± 0.832 | NS |
|  | Circumferential Epicardial Displacement (mm) | Control | 0.7236 ± 0.1652 | - |
|  |  | Aging | 0.327 ± 0.190 | NS |
|  |  | HFD | 0.6997 ± 0.2911 | NS |
|  |  | Exercise | 1.353 ± 0.4450 | NS |
|  | Heart Rate (Beats per minute) | Control | 424.554 ± 26.881 | - |
|  |  | Aging | 452.529 ± 19.063 | NS |
|  |  | HFD | 378.991 ± 33.844 | NS |
|  |  | Exercise | 361.489 ± 25.163 | NS |
|  | Systolic Diameter (mm) | Control | 3.2230 ± 0.1194 | - |
|  |  | Aging | 3.7086 ± 0.1540 | * |
|  |  | HFD | 3.4363 ± 0.1478 | NS |
|  |  | Exercise | 3.1839 ± 0.1356 | NS |
|  | Diastolic Diameter (mm) | Control | 4.4796 ± 0.1179 | - |
|  |  | Aging | 4.6628 ± 0.1668 | NS |
|  |  | HFD | 4.4203 ± 0.1047 | NS |
|  |  | Exercise | 4.4960 ± 0.0290 | NS |
|  | Stroke Volume (uL) | Control | 46.7264 ± 3.6885 | - |
|  |  | Aging | 42.1387 ± 4.0563 | NS |
|  |  | HFD | 39.7614 ± 1.5755 | NS |
|  |  | Exercise | 51.3930 ± 2.9301 | NS |
|  | End Diastolic Volume (uL) | Control | 92.0556 ± 5.8360 | - |
|  |  | Aging | 101.102 ± 8.706 | NS |
|  |  | HFD | 88.8895 ± 4.9769 | NS |
|  |  | Exercise | 92.2830 ± 1.3970 | NS |
|  | Ejection Fraction (%) | Control | 50.8996± 2.9039 | - |
|  |  | Aging | 41.7531 ± 2.5765 | NS (0.06) |
|  |  | HFD | 45.1313 ± 2.9031 | NS |
|  |  | Exercise | 55.8235 ± 3.8250 | NS |
|  | Fractional Shortening (%) | Control | 25.9878 ± 1.8443 | - |
|  |  | Aging | 20.4741 ± 1.4890 | NS (0.07) |
|  |  | HFD | 22.3532 ± 1.6400 | NS |
|  |  | Exercise | 29.2242 ± 2.6713 | NS |
|  | Left Ventricular Posterior wall thickness at systole (mm) | Control | 0.9772 ± 0.0915 | - |
|  |  | Aging | 1.0759 ± 0.0654 | NS |
|  |  | HFD | 0.9749 ± 0.1114 | NS |
|  |  | Exercise | 1.1740 ± 0.1777 | NS |
|  | Left Ventricular Posterior wall thickness at diastole (mm) | Control | 0.5989 ± 0.0188 | - |
|  |  | Aging | 1.1747 ± 0.0897 | * |
|  |  | HFD | 0.7513 ± 0.0381 | * |
|  |  | Exercise | 0.7773 ± 0.0644 | * |
|  | Left Ventricular Mass (mg) | Control | 93.0117 ± 3.9844 | - |
|  |  | Aging | 155.365 ± 18.910 | * |
|  |  | HFD | 108.883 ± 5.183 | * |
|  |  | Exercise | 113.652 ± 9.158 | * |
|  | Cardiac Output (uL/min) | Control | 19.6391 ± 1.6295 | - |
|  |  | Aging | 19.1858 ± 2.3519 | NS |
|  |  | HFD | 15.0788 ± 1.4877 | NS (0.09) |
|  |  | Exercise | 18.7596 ± 2.3648 | NS |
|  | Concentricity (LVPW;d/EDD) | Control | 0.00695 ± 0.00047 | - |
|  |  | Aging | 0.01183 ± 0.00126 | * |
|  |  | HFD | 0.00859 ± 0.00085 | NS (0.09) |
|  |  | Exercise | 0.00845 ± 0.00080 | NS |
|  | Body Weight (g) | Control | 25 ± 1 | - |
|  |  | Aging | 31 ± 2 | * |
|  |  | HFD | 50 ± 2 | * |
|  |  | Exercise | 30 ± 1 | * |
|  | LVPW;d/Body Weight (mm/g) | Control | 0.024 ± 0.003 | - |
|  |  | Aging | 0.031 ± 0.010 | NS |
|  |  | HFD | 0.015 ± 0.003 | * |
|  |  | Exercise | 0.026 ± 0.006 | NS |
| LV Mass/Body Weight (mg/g) | | Control | 3.8 ± 0.6 | - |
|  |  | Aging | 5.0 ± 0.6 | NS |
|  |  | HFD | 2.2 ± 0.4 | * |
|  |  | Exercise | 3.8 ± 0.4 | NS |

*p≤0.05, t-test vs. control

| **Supplemental Table 2: Baseline Echocardiography, Aging v. HFD** | |
| --- | --- |
| **Baseline Measurement** | **P value: T test** |
| Radial Strain | NS |
| Radial Strain Rate | NS |
| Reverse Radial Strain Rate | NS |
| Radial Endocardial Displacement | NS |
| Radial Epicardial Displacement | NS |
| Circumferential Strain | NS |
| Circumferential Strain Rate | NS |
| Reverse Circumferential Strain Rate | NS |
| Circumferential Endocardial Displacement | * |
| Circumferential Epicardial Displacement | NS |
| Heart Rate | NS |
| Systolic Diameter | NS |
| Diastolic Diameter | NS |
| Stroke Volume | NS |
| Ejection Fraction | NS |
| Fractional Shortening | NS |
| LV Posterior wall thickness at systole | NS |
| LV Posterior wall thickness at diastole | NS |
| Cardiac Output | NS |
| End Diastolic Volume | NS |
| LV Mass | NS (0.05) |
| Concentricity | NS (0.07) |
| LVPW;d/Body Weight | * |
| LV Mass/Body Weight | * |

*p≤0.05

| **Supplemental Table 3: Preload Echocardiography** | | | | | | |
| --- | --- | --- | --- | --- | --- | --- |
| **Parameter** | **Group** | **Mean SEM** | **2way ANOVA P value** | | | **Sidak’s**  **p value (v. Baseline)** |
|  |  |  | **Treatment**  **(row)** | **Model**  **(column)** | **R x C**  **(interaction)** |  |
| Radial Strain (%) | Control | 37.4393 ± 3.384 | - | - | - | NS |
|  | Aging | 29.0266 ± 6.7901 | NS | * | NS | NS |
|  | HFD | 32.0100 ± 5.3730 | NS | NS (0.06) | * | * |
|  | Exercise | 39.4434 ± 5.0483 | NS | NS | NS | NS |
| Radial Strain Rate (%/s) | Control | 7.6542 ± 0.6520 | - | - | - | NS |
|  | Aging | 6.5144 ± 1.1807 | NS | * | NS | NS |
|  | HFD | 6.0710 ± 0.7993 | NS | * | * | * |
|  | Exercise | 7.5732 ± 1.0872 | NS | NS | NS | NS |
| Reverse Peak Radial Strain Rate (%/s) | Control | -9.4037 ± 0.6575 | - | - | - | NS |
|  | Aging | -7.6725 ± 1.8877 | NS | NS (0.05) | NS | NS |
|  | HFD | -8.4703 ± 1.4453 | NS (0.08) | NS (0.09) | * | * |
|  | Exercise | -10.1222 ± 1.3269 | NS | NS | NS | NS |
| Radial Endocardial Displacement (mm) | Control | 0.4410 ± 0.0273 | - | - | - | NS |
|  | Aging | 0.449 ± 0.058 | * | NS | NS (0.09) | * |
|  | HFD | 0.3436 ± 0.0410 | NS | NS (0.07) | NS | NS |
|  | Exercise | 0.4200 ± 0.0345 | NS | NS | NS | NS |
| Radial Epicardial Displacement (mm) | Control | 0.2215 ± 0.0221 | - | - | - | * |
|  | Aging | 0.230 ± 0.014 | * | NS | NS | * |
|  | HFD | 0.1505 ± 0.0183 | NS | NS | NS | NS |
|  | Exercise | 0.2214 ± 0.0377 | * | NS | NS | NS |
| Circumferential Strain (%) | Control | -22.0313 ± 1.8650 | - | - | - | NS |
|  | Aging | -18.0627 ± 2.3314 | NS | * | NS | NS |
|  | HFD | -15.3383 ± 1.4555 | NS | * | NS | NS |
|  | Exercise | -20.2099 ± 1.9557 | NS | NS | NS | NS |
| Circumferential Strain Rate (%/s) | Control | -6.6027 ± 0.9709 | - | - | - | NS |
|  | Aging | -5.1624 ± 0.6918 | NS | NS | NS (0.08) | NS |
|  | HFD | -4.0024 ± 0.1851 | NS | * | NS | NS |
|  | Exercise | -5.7053 ± 0.8257 | * | NS | NS | NS |
| Reverse Circumferential Strain Rate (%/s) | Control | 7.2079 ± 0.4085 | - | - | - | NS |
|  | Aging | 6.8969 ± 1.0696 | NS | NS | NS | NS |
|  | HFD | 5.2027 ± 0.1833 | NS | * | NS | NS |
|  | Exercise | 7.4078 ± 1.1059 | NS | NS | NS | NS |
| Circumferential Endocardial Displacement (mm) | Control | 1.051 ± 0.317 | - | - | - | NS (0.05) |
|  | Aging | 1.231 ± 0.511 | NS | NS | * | NS |
|  | HFD | 2.988 ± 0.875 | NS | NS | * | NS |
|  | Exercise | 2.578 ± 1.367 | NS | NS | NS | NS |
| Circumferential Epicardial Displacement (mm) | Control | 0.8417 ± 0.2929 | - | - | - | NS |
|  | Aging | 0.5237 ± 0.1924 | NS | NS | NS | NS |
|  | HFD | 0.7671 ± 0.2475 | NS | NS | NS | NS |
|  | Exercise | 1.156 ± 0.706 | NS | NS | NS | NS |
| Heart Rate (Beats per minute) | Control | 404.373 ± 26.519 | - | - | - | NS |
|  | Aging | 437.931 ± 6.608 | NS | NS | NS | NS |
|  | HFD | 324.375 ± 24.066 | NS (0.06) | NS | NS | NS |
|  | Exercise | 393.134 ± 30.756 | NS | NS | * | NS |
| Systolic Diameter (mm) | Control | 3.3922 ± 0.0858 | - | - | - | * |
|  | Aging | 3.4787 ± 0.2434 | NS | NS | * | NS (0.09) |
|  | HFD | 3.4357 ± 0.1575 | NS | NS | NS (0.08) | NS |
|  | Exercise | 3.4399 ± 0.1056 | * | NS | NS | NS (0.06) |
| Diastolic Diameter (mm) | Control | 4.7945 ± 0.1172 | - | - | - | * |
|  | Aging | 4.6628 ± 0.1668 | * | NS | * | NS |
|  | HFD | 4.7300 ± 0.0933 | * | NS | NS | NS |
|  | Exercise | 4.7915 ± 0.0374 | * | NS | NS | * |
| Stroke Volume (uL) | Control | 60.2813 ± 6.3304 | - | - | - | * |
|  | Aging | 49.3384 ± 4.4362 | * | NS | NS | NS |
|  | HFD | 55.0908 ± 0.9244 | * | NS | NS | * |
|  | Exercise | 58.0951 ± 3.2018 | * | NS | NS | NS |
| End Diastolic Volume (uL) | Control | 107.723 ± 5.998 | - | - | - | * |
|  | Aging | 100.649 ± 9.773 | * | NS | * | NS |
|  | HFD | 104.061 ± 4.734 | * | NS | NS | NS |
|  | Exercise | 107.151 ± 1.956 | * | NS | NS | * |
| Ejection Fraction (%) | Control | 55.3084 ± 3.4800 | - | - | - | NS |
|  | Aging | 49.7409 ± 5.0044 | * | NS | NS | * |
|  | HFD | 53.2088 ± 3.0729 | * | NS | NS | * |
|  | Exercise | 54.2557 ± 3.0489 | NS | NS | NS | NS |
| Fractional Shortening (%) | Control | 29.0529 ± 2.3701 | - | - | - | NS |
|  | Aging | 25.4070 ± 3.0657 | * | NS | NS | * |
|  | HFD | 27.4390 ± 1.9672 | * | NS | NS | * |
|  | Exercise | 28.2194 ± 2.0085 | NS | NS | NS | NS |
| LV Posterior wall thickness at systole (mm) | Control | 1.0464 ± 0.0861 | - | - | - | NS |
|  | Aging | 1.3658 ± 0.1024 | * | NS | * | * |
|  | HFD | 1.0930 ± 0.1982 | NS | NS | NS | NS |
|  | Exercise | 1.0824 ± 0.0531 | NS | NS | NS | NS |
| LV Posterior wall thickness at diastole (mm) | Control | 0.5758 ± 0.0524 | - | - | - | NS |
|  | Aging | 1.0244 ± 0.0386 | NS | * | NS | NS |
|  | HFD | 0.7129 ± 0.0659 | NS | * | NS | NS |
|  | Exercise | 0.6950 ± 0.0209 | NS | * | NS | NS |
| Cardiac Output (uL/min) | Control | 24.6088 ± 3.3729 | - | - | - | NS (0.06) |
|  | Aging | 21.6483 ± 2.0818 | * | NS | NS | NS |
|  | HFD | 17.8844 ± 1.4728 | * | NS | NS | NS |
|  | Exercise | 22.9006 ± 2.3400 | * | NS | NS | NS |

*p≤0.05; Row (treatment): Preload injection vs. measurement; Column (model): Control mice vs. mouse model; Sidak’s comparison: Measurement vs. Baseline measurement

| **Supplemental Table 4: Preload Echocardiography, Aging v. HFD** | | | | | |
| --- | --- | --- | --- | --- | --- |
| **Measurement** | **Model** | **Row** | **Column** | **Interaction** | **Sidak’s** |
| Radial Strain | Aging | NS | NS | NS | NS |
|  | HFD |  |  |  | NS |
| Radial Strain Rate | Aging | NS | NS | NS | NS |
|  | HFD |  |  |  | NS |
| Reverse Radial Strain Rate | Aging | NS | NS | NS | NS |
|  | HFD |  |  |  | NS |
| Radial Endocardial Displacement | Aging | NS (0.08) | NS | NS | NS (0.08) |
|  | HFD |  |  |  | NS |
| Radial Epicardial Displacement | Aging | * | NS (0.09) | * | * |
|  | HFD |  |  |  | NS |
| Circumferential Strain | Aging | * | NS | NS | NS (0.07) |
|  | HFD |  |  |  | NS |
| Circumferential Strain Rate | Aging | NS | NS | NS | NS |
|  | HFD |  |  |  | NS |
| Reverse Circumferential Strain Rate | Aging | NS | NS | NS | NS |
|  | HFD |  |  |  | NS |
| Circumferential Endocardial Displacement | Aging | NS | * | NS | NS |
|  | HFD |  |  |  | NS |
| Circumferential Epicardial Displacement (mm) | Aging | NS | NS | NS | NS |
|  | HFD |  |  |  | NS |
| Heart Rate | Aging | NS | * | NS | NS |
|  | HFD |  |  |  | NS |
| Systolic Diameter | Aging | NS (0.09) | NS | NS | NS (0.09) |
|  | HFD |  |  |  | NS |
| Diastolic Diameter | Aging | * | NS | * | NS |
|  | HFD |  |  |  | * |
| Stroke Volume | Aging | * | NS | NS | NS (0.09) |
|  | HFD |  |  |  | * |
| Ejection Fraction | Aging | * | NS | NS | NS (0.06) |
|  | HFD |  |  |  | NS (0.08) |
| Fractional Shortening | Aging | * | NS | NS | NS (0.06) |
|  | HFD |  |  |  | NS (0.08) |
| LV Posterior wall thickness at systole | Aging | NS (0.09) | NS | NS | NS |
|  | HFD |  |  |  | NS |
| LV Posterior wall thickness at diastole | Aging | NS | * | NS | NS |
|  | HFD |  |  |  | NS |
| Cardiac Output | Aging | NS (0.08) | NS | NS | NS |
|  | HFD |  |  |  | NS |
| End Diastolic Volume | Aging | * | NS | * | NS |
|  | HFD |  |  |  | * |

*p≤0.05; Row (treatment): Preload injection vs. measurement; Column (model): Control mice vs. mouse model; Sidak’s comparison: Measurement vs. Baseline measurement

| **Supplemental Table 5: Dobutamine Echocardiography** | | | | | | |
| --- | --- | --- | --- | --- | --- | --- |
| **Parameter** | **Group** | **Mean SEM** | **2way ANOVA P value** | | | **Sidak’s**  **p value (v. Baseline)** |
|  |  |  | **Row**  **(treatment)** | **Column**  **(models)** | **R x C**  **(interaction)** |  |
| Radial Strain (%) | Control | 57.986 ± 4.090 | - | - | - | * |
|  | Aging | 29.298 ± 1.166 | * | * | NS | NS (0.07) |
|  | HFD | 37.666 ± 4.796 | * | * | * | * |
|  | Exercise | 63.610 ± 6.190 | * | NS | NS | * |
| Radial Strain Rate (%/s) | Control | 14.0525 ± 1.1774 | - | - | - | * |
|  | Aging | 8.4144 ± 0.6570 | * | * | NS | * |
|  | HFD | 9.6576 ± 1.4472 | * | * | NS | * |
|  | Exercise | 17.0143 ± 1.0251 | * | NS | NS (0.06) | * |
| Reverse Peak Radial Strain Rate (%/s) | Control | -14.6109 ± 1.0608 | - | - | - | * |
|  | Aging | -8.8791 ± 0.6352 | * | * | NS (0.07) | NS |
|  | HFD | -10.6174 ± 0.6013 | * | * | * | * |
|  | Exercise | -18.1403 ± 1.6338 | * | NS | * | * |
| Radial Endocardial Displacement (mm) | Control | 0.6344 ± 0.0169 | - | - | - | * |
|  | Aging | 0.517 ± 0.021 | * | * | NS | * |
|  | HFD | 0.5768 ± 0.0419 | * | NS (0.06) | NS | * |
|  | Exercise | 0.6787 ± 0.0252 | * | NS | NS | * |
| Radial Epicardial Displacement (mm) | Control | 0.2347 ± 0.0151 | - | - | - | * |
|  | Aging | 0.2564 ± 0.0213 | * | NS | NS | * |
|  | HFD | 0.2785 ± 0.0115 | * | NS | * | * |
|  | Exercise | 0.2162 ± 0.0194 | * | NS | NS | NS |
| Circumferential Strain (%) | Control | -33.1930 ± 1.1870 | - | - | - | * |
|  | Aging | -24.5169 ± 2.2878 | * | * | NS | * |
|  | HFD | -25.4276 ± 2.8411 | * | * | NS | * |
|  | Exercise | -35.9170 ± 0.9975 | * | NS | NS (0.09) | * |
| Circumferential Strain Rate (%/s) | Control | -13.8040 ± 0.7343 | - | - | - | * |
|  | Aging | -10.2836 ± 1.9989 | * | NS (0.05) | NS | * |
|  | HFD | -10.6721 ± 0.8957 | * | * | NS | * |
|  | Exercise | -18.5150 ± 1.3053 | * | NS (0.05) | * | * |
| Reverse Circumferential Strain Rate (%/s) | Control | 14.2197 ± 1.0395 | - | - | - | * |
|  | Aging | 9.5994 ± 1.7605 | * | * | NS | NS (0.05) |
|  | HFD | 11.8862 ± 1.7230 | * | NS (0.05) | NS | * |
|  | Exercise | 16.2244 ± 1.1516 | * | NS | NS | * |
| Circumferential Endocardial Displacement (mm) | Control | 4.099 ± 1.191 | - | - | - | NS (0.06) |
|  | Aging | 0.5222 ± 0.2649 | NS | * | NS | NS |
|  | HFD | 2.910 ± 1.543 | NS (0.08) | NS | NS | NS |
|  | Exercise | 4.209 ± 1.385 | NS | NS | NS | NS |
| Circumferential Epicardial Displacement (mm) | Control | 0.712 ± 0.104 | - | - | - | NS |
|  | Aging | 1.150 ± 0.165 | * | NS | * | * |
|  | HFD | 1.327 ± 0.215 | NS | NS | NS | NS (0.09) |
|  | Exercise | 1.748 ± 0.530 | NS | * | NS | NS |
| Heart Rate (Beats per minute) | Control | 543.112 ± 14.748 | - | - | - | * |
|  | Aging | 538.138 ± 16.540 | * | NS | NS | * |
|  | HFD | 534.773 ± 37.622 | * | NS | NS | * |
|  | Exercise | 595.169 ± 9.649 | * | NS | * | * |
| Systolic Diameter (mm) | Control | 2.0363 ± 0.1192 | - | - | - | * |
|  | Aging | 2.4930 ± 0.4818 | * | NS (0.07) | NS | * |
|  | HFD | 2.0103 ± 0.2129 | * | NS | NS | * |
|  | Exercise | 1.7013 ± 0.1464 | * | NS | NS | * |
| Diastolic Diameter (mm) | Control | 3.8098 ± 0.0965 | - | - | - | * |
|  | Aging | 4.0423 ± 0.4137 | * | NS | NS | NS (0.05) |
|  | HFD | 3.8594 ± 0.1852 | * | NS | NS | NS (0.06) |
|  | Exercise | 3.7102 ± 0.1235 | * | NS | NS | * |
| Stroke Volume (uL) | Control | 48.8891 ± 2.3377 | - | - | - | NS |
|  | Aging | 48.7086 ± 6.3833 | NS | NS | NS | NS |
|  | HFD | 51.2729 ± 4.7182 | * | NS | NS | NS (0.05) |
|  | Exercise | 50.0450 ± 2.7815 | NS | NS | NS | NS |
| End Diastolic Volume (uL) | Control | 62.7620 ± 3.7670 | - | - | - | * |
|  | Aging | 75.2524 ± 17.6226 | * | NS | NS | NS (0.05) |
|  | HFD | 64.9851 ± 6.9932 | * | NS | NS | NS (0.07) |
|  | Exercise | 58.8475 ± 4.6796 | * | NS | NS | * |
| Ejection Fraction (%) | Control | 78.3761 ± 2.2420 | - | - | - | * |
|  | Aging | 69.7912 ± 7.3049 | * | * | NS | * |
|  | HFD | 79.5944 ± 3.6674 | * | NS | NS | * |
|  | Exercise | 85.5271 ± 2.1192 | * | NS | NS | * |
| Fractional Shortening (%) | Control | 46.7353 ± 2.1512 | - | - | - | * |
|  | Aging | 40.0331 ± 5.6077 | * | NS (0.06) | NS | * |
|  | HFD | 48.1777 ± 3.7745 | * | NS | NS | * |
|  | Exercise | 54.3770 ± 2.4095 | * | NS (0.06) | NS | * |
| LV Posterior wall thickness at systole (mm) | Control | 1.3682 ± 0.0648 | - | - | - | * |
|  | Aging | 1.6550 ± 0.0476 | * | NS (0.08) | NS | * |
|  | HFD | 1.8205 ± 0.1299 | * | NS (0.05) | * | * |
|  | Exercise | 1.6200 ± 0.0669 | * | NS | NS | * |
| LV Posterior wall thickness at diastole (mm) | Control | 0.7943 ± 0.0570 | - | - | - | NS |
|  | Aging | 1.1747 ± 0.0897 | * | * | NS | NS |
|  | HFD | 1.2353 ± 0.1361 | * | * | NS (0.08) | * |
|  | Exercise | 0.8896 ± 0.0270 | * | * | NS | NS |
| Cardiac Output (uL/min) | Control | 26.7122 ± 1.8541 | - | - | - | * |
|  | Aging | 25.9111 ± 2.8179 | * | NS | NS | * |
|  | HFD | 27.6773 ± 3.7352 | * | NS | NS (0.08) | * |
|  | Exercise | 29.8539 ± 2.0666 | * | NS | NS | * |

*p≤0.05; Row (treatment): Preload injection vs. measurement; Column (model): Control mice vs. mouse model; Sidak’s comparison: Measurement vs. Baseline measurement

| **Supplemental Table 6: Dobutamine Echocardiography, Aging v. HFD** | | | | | |
| --- | --- | --- | --- | --- | --- |
| **Measurement** | **Model** | **Row** | **Column** | **Interaction** | **Sidak’s** |
| Radial Strain | Aging | * | NS | NS | NS (0.09) |
|  | HFD |  |  |  | NS (0.08) |
| Radial Strain Rate | Aging | * | NS | NS | * |
|  | HFD |  |  |  | * |
| Reverse Radial Strain Rate | Aging | * | NS | NS | NS |
|  | HFD |  |  |  | * |
| Radial Endocardial Displacement | Aging | * | NS | NS | * |
|  | HFD |  |  |  | * |
| Radial Epicardial Displacement | Aging | * | NS | NS | * |
|  | HFD |  |  |  | * |
| Circumferential Strain | Aging | * | NS | NS | * |
|  | HFD |  |  |  | * |
| Circumferential Strain Rate | Aging | * | NS | NS | * |
|  | HFD |  |  |  | * |
| Reverse Circumferential Strain Rate | Aging | * | NS | NS | * |
|  | HFD |  |  |  | * |
| Circumferential Endocardial Displacement | Aging | NS | NS (0.08) | NS | NS |
|  | HFD |  |  |  | NS |
| Circumferential Epicardial Displacement (mm) | Aging | * | * | NS | NS |
|  | HFD |  |  |  | NS |
| Heart Rate | Aging | * | NS | NS | NS (0.05) |
|  | HFD |  |  |  | * |
| Systolic Diameter | Aging | * | NS | NS | * |
|  | HFD |  |  |  | * |
| Diastolic Diameter | Aging | * | NS | NS | NS |
|  | HFD |  |  |  | NS |
| Stroke Volume | Aging | * | NS | NS | NS |
|  | HFD |  |  |  | NS (0.08) |
| Ejection Fraction | Aging | * | NS | NS | * |
|  | HFD |  |  |  | * |
| Fractional Shortening | Aging | * | NS | NS | * |
|  | HFD |  |  |  | * |
| LV Posterior wall thickness at systole | Aging | * | NS | NS | * |
|  | HFD |  |  |  | * |
| LV Posterior wall thickness at diastole | Aging | * | NS | NS | NS |
|  | HFD |  |  |  | NS (0.06) |
| Cardiac Output | Aging | * | NS | NS | NS |
|  | HFD |  |  |  | * |
| End Diastolic Volume | Aging | * | NS | NS | NS |
|  | HFD |  |  |  | NS |

*p≤0.05; Row (treatment): Preload injection vs. measurement; Column (model): Aging mice vs. HFD model; Sidak’s comparison: Measurement vs. Baseline measurement

| **Supplemental Table 7: 8 Week High Intensity Interval Training (HIIT) Exercise Protocol** | | | | | |
| --- | --- | --- | --- | --- | --- |
| **Weeks** | **Incline** | **Total Daily Time** | **Number of Sets** | **Fast Pace (m/min)** | **Recovery Pace (m/min)** |
| 1 | 5 | 40 | 8 | 18 | 10 |
| 2 | 10 | 50 | 10 | 20 | 10 |
| 3 | 15 | 60 | 12 | 22 | 11 |
| 4 | 20 | 70 | 14 | 24 | 11 |
| 5 | 20 | 80 | 16 | 26 | 12 |
| 6 | 20 | 80 | 16 | 28 | 12 |
| 7 | 20 | 80 | 16 | 30 | 13 |
| 8 | 20 | 80 | 16 | 32 | 13 |
